## Supplementary Materials for "Viscoelastic characteristics of the canine cranial cruciate ligament complex at slow strain rates"

Tables

**Table S1**: P-values obtained from Bonferroni post-hoc statistical analysis on ligament lengths at different planes.

| **Cranial and Caudal Planes** | **Cranial and Medial Planes** | **Cranial and Lateral Planes** | **Caudal and Medial Planes** | **Caudal and Lateral Planes** | **Medial and Lateral Planes** |
| --- | --- | --- | --- | --- | --- |
| $6.74\times{10}^{-7}$ | $9.96\times{10}^{-3}$ | $2.25\times{10}^{-2}$ | $1.49\times{10}^{-5}$ | $1.41\times{10}^{-4}$ | $9.07\times{10}^{-1}$ |

Figures

| 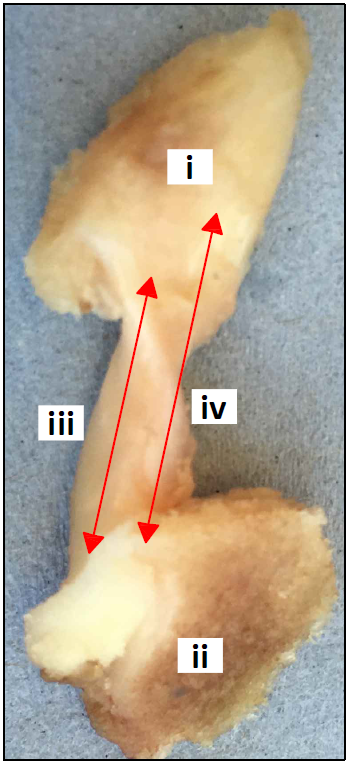  **(a)** | 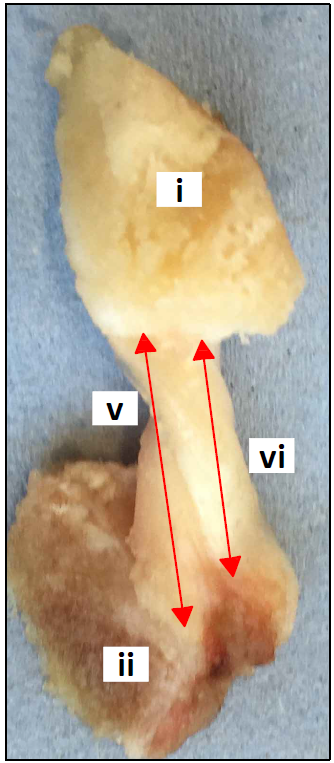  **(b)** |
| --- | --- |

Fig. S1. The cranial cruciate ligament (CCL) from a right canine stifle joint from the (a) cranial, and (b) caudal views of the ligament. The length of the CCL, attached to the (i) femur and (ii) tibia was measured at four different planes: (iii) lateral, (iv) cranial, (v) medial and (vi) caudal.

| 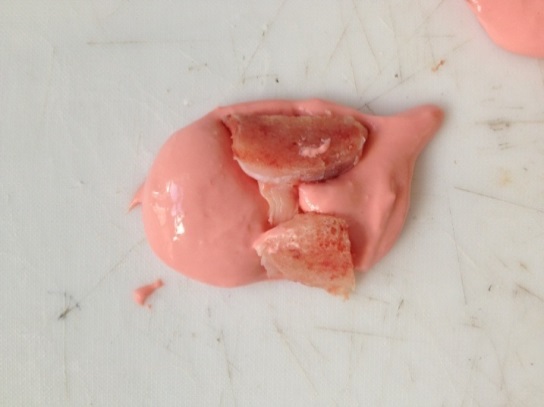  **(a)**  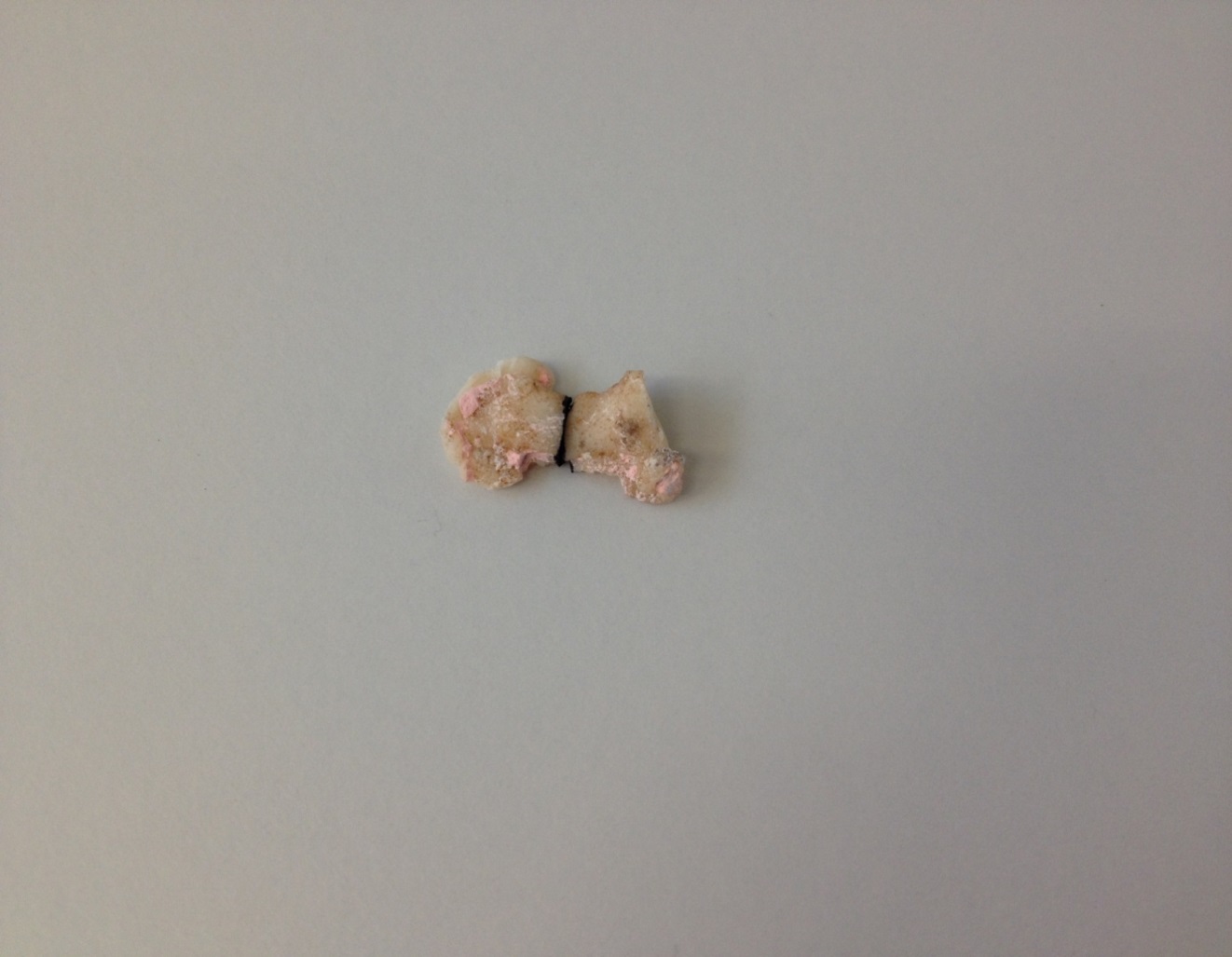  **(c)** | 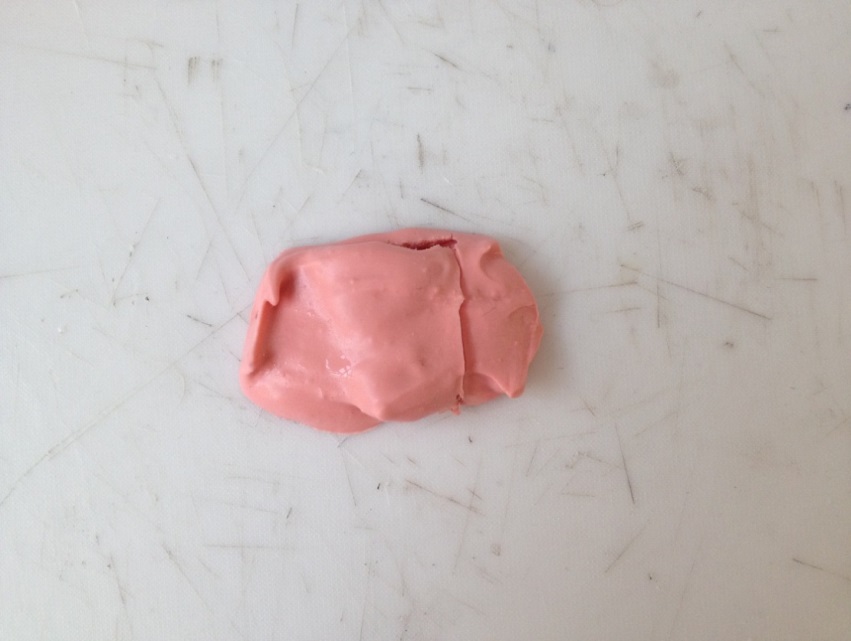  **(b)**  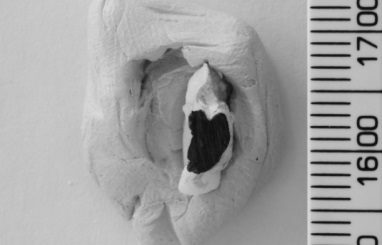  **(d)** |
| --- | --- |

Fig. S2. The middle cross-sectional area (CSA) of the cranial cruciate ligament (CCL) was estimated by firstly creating an alginate paste and (a) place the paste around the CCL until it was set. (b) Subsequently, the CCL was removed from the set paste leaving behind an alginate mould for the ligament. (c) Poly methyl methacrylate (PMMA) was injected into the mould to create replica model of the CCL. The replica was cut into two in the middle and (d) the surface of the replica showing middle CSA was coloured in black paint. The black surface area of the replica was then measured using ImageJ (a public domain Java image processing program).

Stress (MPa)

| Strain (mm/mm) | CCL1_Desc |
| --- | --- |
| 0 | 0 |
| 0.0025 | 0.000910528 |
| 0.005 | 0.00188246 |
| 0.0075 | 0.00291994 |
| 0.01 | 0.00402739 |
| 0.0125 | 0.00520952 |
| 0.015 | 0.00647138 |
| 0.0175 | 0.00781833 |
| 0.02 | 0.00925612 |
| 0.0225 | 0.0107909 |
| 0.025 | 0.0124291 |
| 0.0275 | 0.0141779 |
| 0.03 | 0.0160446 |
| 0.0325 | 0.0180371 |
| 0.035 | 0.0201641 |
| 0.0375 | 0.0224344 |
| 0.04 | 0.0248579 |
| 0.0425 | 0.0274449 |
| 0.045 | 0.0302063 |
| 0.0475 | 0.0331539 |
| 0.05 | 0.0363003 |
| 0.0525 | 0.0396589 |
| 0.055 | 0.043244 |
| 0.0575 | 0.0470709 |
| 0.06 | 0.0511558 |
| 0.0625 | 0.0555163 |
| 0.065 | 0.0601708 |
| 0.0675 | 0.0651392 |
| 0.07 | 0.0704427 |
| 0.0725 | 0.0761038 |
| 0.075 | 0.0821468 |
| 0.0775 | 0.0885972 |
| 0.08 | 0.0954827 |
| 0.0825 | 0.102833 |
| 0.085 | 0.110678 |
| 0.0875 | 0.119053 |
| 0.09 | 0.127992 |
| 0.0925 | 0.137534 |
| 0.095 | 0.14772 |
| 0.0975 | 0.158593 |
| 0.1 | 0.170199 |
| 0.1025 | 0.182587 |
| 0.105 | 0.195811 |
| 0.1075 | 0.209927 |
| 0.11 | 0.224995 |
| 0.1125 | 0.241079 |
| 0.115 | 0.258248 |
| 0.1175 | 0.276575 |
| 0.12 | 0.296137 |
| 0.1225 | 0.317019 |
| 0.125 | 0.339309 |
| 0.1275 | 0.363103 |
| 0.13 | 0.388501 |
| 0.1325 | 0.415611 |
| 0.135 | 0.44455 |
| 0.1375 | 0.475441 |
| 0.14 | 0.508415 |
| 0.1425 | 0.543613 |
| 0.145 | 0.581185 |
| 0.1475 | 0.62129 |
| 0.15 | 0.6641 |
| 0.1525 | 0.709797 |
| 0.155 | 0.758576 |
| 0.1575 | 0.810644 |

| Strain (mm/mm) | CCL1_Asc |
| --- | --- |
| 0 | 0 |
| 0.0025 | 0.00383344 |
| 0.005 | 0.00811158 |
| 0.0075 | 0.012886 |
| 0.01 | 0.0182143 |
| 0.0125 | 0.0241608 |
| 0.015 | 0.030797 |
| 0.0175 | 0.0382031 |
| 0.02 | 0.0464684 |
| 0.0225 | 0.0556925 |
| 0.025 | 0.0659866 |
| 0.0275 | 0.077475 |
| 0.03 | 0.0902961 |
| 0.0325 | 0.104604 |
| 0.035 | 0.120573 |
| 0.0375 | 0.138393 |
| 0.04 | 0.158282 |
| 0.0425 | 0.180477 |
| 0.045 | 0.205247 |
| 0.0475 | 0.23289 |
| 0.05 | 0.263741 |
| 0.0525 | 0.29817 |
| 0.055 | 0.336593 |
| 0.0575 | 0.379474 |
| 0.06 | 0.427329 |
| 0.0625 | 0.480735 |
| 0.065 | 0.540337 |
| 0.0675 | 0.606853 |

Stress (MPa)

| Strain (mm/mm) | CCL2_Desc |
| --- | --- |
| 0 | 0 |
| 0.0025 | 0.00379406 |
| 0.005 | 0.00874799 |
| 0.0075 | 0.0152164 |
| 0.01 | 0.0236622 |
| 0.0125 | 0.0346899 |
| 0.015 | 0.049089 |
| 0.0175 | 0.0678899 |
| 0.02 | 0.0924383 |
| 0.0225 | 0.124491 |
| 0.025 | 0.166343 |
| 0.0275 | 0.22099 |
| 0.03 | 0.292342 |
| 0.0325 | 0.385507 |
| 0.035 | 0.507154 |
| 0.0375 | 0.665988 |

| Strain (mm/mm) | CCL2_Asc |
| --- | --- |
| 0 | 0 |
| 0.0025 | 0.00204074 |
| 0.005 | 0.00440673 |
| 0.0075 | 0.00714981 |
| 0.01 | 0.0103301 |
| 0.0125 | 0.0140172 |
| 0.015 | 0.018292 |
| 0.0175 | 0.0232481 |
| 0.02 | 0.0289941 |
| 0.0225 | 0.0356559 |
| 0.025 | 0.0433794 |
| 0.0275 | 0.0523339 |
| 0.03 | 0.0627155 |
| 0.0325 | 0.0747517 |
| 0.035 | 0.0887063 |
| 0.0375 | 0.104885 |
| 0.04 | 0.123642 |
| 0.0425 | 0.145389 |
| 0.045 | 0.170601 |
| 0.0475 | 0.199832 |
| 0.05 | 0.233722 |
| 0.0525 | 0.273013 |
| 0.055 | 0.318566 |
| 0.0575 | 0.37138 |
| 0.06 | 0.43261 |
| 0.0625 | 0.5036 |
| 0.065 | 0.585904 |
| 0.0675 | 0.681325 |

Stress (MPa)

| Strain (mm/mm) | CCL3_Desc |
| --- | --- |
| 0 | 0 |
| 0.0025 | 0.00194154 |
| 0.005 | 0.00414699 |
| 0.0075 | 0.00665222 |
| 0.01 | 0.00949797 |
| 0.0125 | 0.0127305 |
| 0.015 | 0.0164025 |
| 0.0175 | 0.0205736 |
| 0.02 | 0.0253116 |
| 0.0225 | 0.0306936 |
| 0.025 | 0.0368072 |
| 0.0275 | 0.0437519 |
| 0.03 | 0.0516404 |
| 0.0325 | 0.0606013 |
| 0.035 | 0.0707801 |
| 0.0375 | 0.0823425 |
| 0.04 | 0.0954765 |
| 0.0425 | 0.110396 |
| 0.045 | 0.127343 |
| 0.0475 | 0.146594 |
| 0.05 | 0.168461 |
| 0.0525 | 0.193301 |
| 0.055 | 0.221517 |
| 0.0575 | 0.253569 |
| 0.06 | 0.289977 |
| 0.0625 | 0.331334 |
| 0.065 | 0.378313 |
| 0.0675 | 0.431677 |
| 0.07 | 0.492295 |
| 0.0725 | 0.561152 |
| 0.075 | 0.639369 |
| 0.0775 | 0.728218 |
| 0.08 | 0.829143 |
| 0.0825 | 0.943787 |

| Strain (mm/mm) | CCL3_Asc |
| --- | --- |
| 0 | 0 |
| 0.0025 | 0.00195749 |
| 0.005 | 0.00423884 |
| 0.0075 | 0.00689764 |
| 0.01 | 0.00999634 |
| 0.0125 | 0.0136077 |
| 0.015 | 0.0178166 |
| 0.0175 | 0.0227218 |
| 0.02 | 0.0284386 |
| 0.0225 | 0.0351013 |
| 0.025 | 0.0428662 |
| 0.0275 | 0.0519159 |
| 0.03 | 0.0624629 |
| 0.0325 | 0.0747548 |
| 0.035 | 0.0890804 |
| 0.0375 | 0.105776 |
| 0.04 | 0.125234 |
| 0.0425 | 0.147912 |
| 0.045 | 0.174341 |
| 0.0475 | 0.205143 |
| 0.05 | 0.241041 |
| 0.0525 | 0.282879 |
| 0.055 | 0.331639 |

Stress (MPa)

| Strain (mm/mm) | CCL4_Desc |
| --- | --- |
| 0 | 0 |
| 0.0025 | 0.00192952 |
| 0.005 | 0.00416489 |
| 0.0075 | 0.00675456 |
| 0.01 | 0.00975472 |
| 0.0125 | 0.0132304 |
| 0.015 | 0.017257 |
| 0.0175 | 0.0219219 |
| 0.02 | 0.0273261 |
| 0.0225 | 0.033587 |
| 0.025 | 0.0408402 |
| 0.0275 | 0.0492431 |
| 0.03 | 0.058978 |
| 0.0325 | 0.0702558 |
| 0.035 | 0.0833212 |
| 0.0375 | 0.0984576 |
| 0.04 | 0.115993 |
| 0.0425 | 0.136308 |
| 0.045 | 0.159843 |
| 0.0475 | 0.187109 |
| 0.05 | 0.218696 |
| 0.0525 | 0.25529 |
| 0.055 | 0.297684 |
| 0.0575 | 0.346798 |
| 0.06 | 0.403697 |
| 0.0625 | 0.469615 |
| 0.065 | 0.545981 |
| 0.0675 | 0.634451 |

| Strain (mm/mm) | CCL4_Asc |
| --- | --- |
| 0 | 0 |
| 0.0025 | 0.00138486 |
| 0.005 | 0.00294009 |
| 0.0075 | 0.00468663 |
| 0.01 | 0.00664801 |
| 0.0125 | 0.00885069 |
| 0.015 | 0.0113243 |
| 0.0175 | 0.0141022 |
| 0.02 | 0.0172219 |
| 0.0225 | 0.0207253 |
| 0.025 | 0.0246597 |
| 0.0275 | 0.0290781 |
| 0.03 | 0.03404 |
| 0.0325 | 0.0396123 |
| 0.035 | 0.0458701 |
| 0.0375 | 0.0528977 |
| 0.04 | 0.0607897 |
| 0.0425 | 0.0696527 |
| 0.045 | 0.0796058 |
| 0.0475 | 0.0907834 |
| 0.05 | 0.103336 |
| 0.0525 | 0.117433 |
| 0.055 | 0.133264 |
| 0.0575 | 0.151042 |
| 0.06 | 0.171007 |
| 0.0625 | 0.193429 |
| 0.065 | 0.218608 |
| 0.0675 | 0.246885 |
| 0.07 | 0.278641 |
| 0.0725 | 0.314303 |
| 0.075 | 0.354351 |
| 0.0775 | 0.399327 |
| 0.08 | 0.449835 |
| 0.0825 | 0.506556 |
| 0.085 | 0.570255 |
| 0.0875 | 0.64179 |
| 0.09 | 0.722125 |

Stress (MPa)

| Strain (mm/mm) | CCL5_Desc |
| --- | --- |
| 0 | 0 |
| 0.0025 | 0.00130963 |
| 0.005 | 0.00275587 |
| 0.0075 | 0.00435298 |
| 0.01 | 0.00611668 |
| 0.0125 | 0.00806437 |
| 0.015 | 0.0102152 |
| 0.0175 | 0.0125904 |
| 0.02 | 0.0152134 |
| 0.0225 | 0.01811 |
| 0.025 | 0.0213087 |
| 0.0275 | 0.0248412 |
| 0.03 | 0.0287421 |
| 0.0325 | 0.0330499 |
| 0.035 | 0.037807 |
| 0.0375 | 0.0430604 |
| 0.04 | 0.0488618 |
| 0.0425 | 0.0552684 |
| 0.045 | 0.0623433 |
| 0.0475 | 0.0701561 |
| 0.05 | 0.078784 |
| 0.0525 | 0.0883118 |
| 0.055 | 0.0988336 |
| 0.0575 | 0.110453 |
| 0.06 | 0.123284 |
| 0.0625 | 0.137454 |
| 0.065 | 0.153102 |
| 0.0675 | 0.170382 |
| 0.07 | 0.189465 |
| 0.0725 | 0.210538 |
| 0.075 | 0.23381 |
| 0.0775 | 0.259509 |
| 0.08 | 0.287889 |
| 0.0825 | 0.319229 |
| 0.085 | 0.353838 |
| 0.0875 | 0.392058 |
| 0.09 | 0.434264 |
| 0.0925 | 0.480873 |
| 0.095 | 0.532344 |
| 0.0975 | 0.589185 |
| 0.1 | 0.651954 |
| 0.1025 | 0.721271 |
| 0.105 | 0.797819 |

| Strain (mm/mm) | CCL5_Asc |
| --- | --- |
| 0 | 0 |
| 0.0025 | 0.00124152 |
| 0.005 | 0.00262827 |
| 0.0075 | 0.00417725 |
| 0.01 | 0.00590743 |
| 0.0125 | 0.00784001 |
| 0.015 | 0.00999867 |
| 0.0175 | 0.0124099 |
| 0.02 | 0.0151031 |
| 0.0225 | 0.0181114 |
| 0.025 | 0.0214716 |
| 0.0275 | 0.0252249 |
| 0.03 | 0.0294173 |
| 0.0325 | 0.0341001 |
| 0.035 | 0.0393308 |
| 0.0375 | 0.0451733 |
| 0.04 | 0.0516992 |
| 0.0425 | 0.0589886 |
| 0.045 | 0.0671308 |
| 0.0475 | 0.0762254 |
| 0.05 | 0.0863839 |
| 0.0525 | 0.0977308 |
| 0.055 | 0.110405 |
| 0.0575 | 0.124562 |
| 0.06 | 0.140375 |
| 0.0625 | 0.158038 |
| 0.065 | 0.177767 |
| 0.0675 | 0.199804 |
| 0.07 | 0.224419 |
| 0.0725 | 0.251913 |
| 0.075 | 0.282624 |
| 0.0775 | 0.316928 |
| 0.08 | 0.355244 |
| 0.0825 | 0.398043 |
| 0.085 | 0.445848 |
| 0.0875 | 0.499245 |
| 0.09 | 0.55889 |
| 0.0925 | 0.625511 |

Stress (MPa)

| Strain (mm/mm) | S6_Desc |
| --- | --- |
| 0 | 0 |
| 0.0025 | 0.00137659 |
| 0.005 | 0.00293312 |
| 0.0075 | 0.00469311 |
| 0.01 | 0.00668316 |
| 0.0125 | 0.00893334 |
| 0.015 | 0.0114777 |
| 0.0175 | 0.0143546 |
| 0.02 | 0.0176075 |
| 0.0225 | 0.0212857 |
| 0.025 | 0.0254446 |
| 0.0275 | 0.0301472 |
| 0.03 | 0.0354645 |
| 0.0325 | 0.0414769 |
| 0.035 | 0.0482752 |
| 0.0375 | 0.0559621 |
| 0.04 | 0.0646538 |
| 0.0425 | 0.0744817 |
| 0.045 | 0.0855942 |
| 0.0475 | 0.0981593 |
| 0.05 | 0.112367 |
| 0.0525 | 0.128432 |
| 0.055 | 0.146596 |
| 0.0575 | 0.167135 |
| 0.06 | 0.190359 |
| 0.0625 | 0.216618 |
| 0.065 | 0.24631 |
| 0.0675 | 0.279884 |
| 0.07 | 0.317845 |
| 0.0725 | 0.360769 |
| 0.075 | 0.409304 |
| 0.0775 | 0.464183 |
| 0.08 | 0.526236 |
| 0.0825 | 0.5964 |
| 0.085 | 0.675736 |

| Strain (mm/mm) | CCL6_Asc |
| --- | --- |
| 0 | 0 |
| 0.0025 | 0.00150435 |
| 0.005 | 0.00320461 |
| 0.0075 | 0.00512631 |
| 0.01 | 0.00729829 |
| 0.0125 | 0.00975313 |
| 0.015 | 0.0125277 |
| 0.0175 | 0.0156636 |
| 0.02 | 0.0192079 |
| 0.0225 | 0.0232138 |
| 0.025 | 0.0277414 |
| 0.0275 | 0.0328586 |
| 0.03 | 0.0386423 |
| 0.0325 | 0.0451792 |
| 0.035 | 0.0525675 |
| 0.0375 | 0.060918 |
| 0.04 | 0.070356 |
| 0.0425 | 0.0810232 |
| 0.045 | 0.0930796 |
| 0.0475 | 0.106706 |
| 0.05 | 0.122107 |
| 0.0525 | 0.139515 |
| 0.055 | 0.159189 |
| 0.0575 | 0.181425 |
| 0.06 | 0.206557 |
| 0.0625 | 0.234963 |
| 0.065 | 0.267067 |
| 0.0675 | 0.303353 |
| 0.07 | 0.344365 |
| 0.0725 | 0.390718 |
| 0.075 | 0.443107 |

**Fig. S3.** The stress-strain characteristics of individual canine cranial cruciate ligaments (CCL) during the ascending (Asc) and descending (Desc) tests at 0.1%/min strain rate.

Stress (MPa)

| Strain (mm/mm) | CCL1_Desc |
| --- | --- |
| 0 | 0 |
| 0.0025 | 0.000904718 |
| 0.005 | 0.00187174 |
| 0.0075 | 0.00290535 |
| 0.01 | 0.00401014 |
| 0.0125 | 0.00519102 |
| 0.015 | 0.00645321 |
| 0.0175 | 0.00780232 |
| 0.02 | 0.00924434 |
| 0.0225 | 0.0107857 |
| 0.025 | 0.0124331 |
| 0.0275 | 0.014194 |
| 0.03 | 0.0160762 |
| 0.0325 | 0.018088 |
| 0.035 | 0.0202383 |
| 0.0375 | 0.0225367 |
| 0.04 | 0.0249934 |
| 0.0425 | 0.0276193 |
| 0.045 | 0.030426 |
| 0.0475 | 0.033426 |
| 0.05 | 0.0366325 |
| 0.0525 | 0.0400599 |
| 0.055 | 0.0437233 |
| 0.0575 | 0.047639 |
| 0.06 | 0.0518244 |
| 0.0625 | 0.0562979 |
| 0.065 | 0.0610795 |
| 0.0675 | 0.0661905 |
| 0.07 | 0.0716533 |
| 0.0725 | 0.0774924 |
| 0.075 | 0.0837335 |
| 0.0775 | 0.0904045 |
| 0.08 | 0.0975348 |
| 0.0825 | 0.105156 |
| 0.085 | 0.113302 |
| 0.0875 | 0.12201 |
| 0.09 | 0.131316 |
| 0.0925 | 0.141264 |
| 0.095 | 0.151897 |
| 0.0975 | 0.163262 |
| 0.1 | 0.175409 |
| 0.1025 | 0.188393 |
| 0.105 | 0.202272 |
| 0.1075 | 0.217106 |
| 0.11 | 0.232961 |
| 0.1125 | 0.249909 |
| 0.115 | 0.268023 |
| 0.1175 | 0.287385 |
| 0.12 | 0.30808 |
| 0.1225 | 0.330201 |
| 0.125 | 0.353844 |
| 0.1275 | 0.379116 |
| 0.13 | 0.406128 |
| 0.1325 | 0.435001 |
| 0.135 | 0.465861 |
| 0.1375 | 0.498847 |
| 0.14 | 0.534105 |
| 0.1425 | 0.57179 |
| 0.145 | 0.61207 |
| 0.1475 | 0.655125 |
| 0.15 | 0.701144 |
| 0.1525 | 0.750332 |
| 0.155 | 0.802908 |

| Strain (mm/mm) | CCL1_Asc |
| --- | --- |
| 0 | 0 |
| 0.0025 | 0.00346459 |
| 0.005 | 0.00738391 |
| 0.0075 | 0.0118176 |
| 0.01 | 0.0168333 |
| 0.0125 | 0.0225073 |
| 0.015 | 0.0289259 |
| 0.0175 | 0.036187 |
| 0.02 | 0.0444012 |
| 0.0225 | 0.0536934 |
| 0.025 | 0.0642053 |
| 0.0275 | 0.0760968 |
| 0.03 | 0.0895491 |
| 0.0325 | 0.104767 |
| 0.035 | 0.121982 |
| 0.0375 | 0.141457 |
| 0.04 | 0.163488 |
| 0.0425 | 0.18841 |
| 0.045 | 0.216604 |
| 0.0475 | 0.248497 |
| 0.05 | 0.284577 |
| 0.0525 | 0.325393 |
| 0.055 | 0.371565 |
| 0.0575 | 0.423797 |
| 0.06 | 0.482885 |
| 0.0625 | 0.549728 |
| 0.065 | 0.625345 |

Stress (MPa)

| Strain (mm/mm) | CCL2_Desc |
| --- | --- |
| 0 | 0 |
| 0.0025 | 0.00351959 |
| 0.005 | 0.00813883 |
| 0.0075 | 0.0142013 |
| 0.01 | 0.0221579 |
| 0.0125 | 0.0326004 |
| 0.015 | 0.0463056 |
| 0.0175 | 0.0642928 |
| 0.02 | 0.0878999 |
| 0.0225 | 0.118883 |
| 0.025 | 0.159546 |
| 0.0275 | 0.212914 |
| 0.03 | 0.282955 |
| 0.0325 | 0.374881 |
| 0.035 | 0.495527 |
| 0.0375 | 0.653868 |

| Strain (mm/mm) | CCL2_Asc |
| --- | --- |
| 0 | 0 |
| 0.0025 | 0.00221552 |
| 0.005 | 0.00486882 |
| 0.0075 | 0.00804639 |
| 0.01 | 0.0118518 |
| 0.0125 | 0.0164092 |
| 0.015 | 0.0218671 |
| 0.0175 | 0.0284034 |
| 0.02 | 0.0362313 |
| 0.0225 | 0.0456059 |
| 0.025 | 0.0568328 |
| 0.0275 | 0.0702782 |
| 0.03 | 0.0863803 |
| 0.0325 | 0.105664 |
| 0.035 | 0.128758 |
| 0.0375 | 0.156415 |
| 0.04 | 0.189538 |
| 0.0425 | 0.229205 |
| 0.045 | 0.27671 |
| 0.0475 | 0.333602 |
| 0.05 | 0.401735 |
| 0.0525 | 0.483331 |
| 0.055 | 0.581049 |
| 0.0575 | 0.698077 |

Stress (MPa)

| Strain (mm/mm) | CCL3_Desc |
| --- | --- |
| 0 | 0 |
| 0.0025 | 0.00184447 |
| 0.005 | 0.00395366 |
| 0.0075 | 0.0063656 |
| 0.01 | 0.00912371 |
| 0.0125 | 0.0122777 |
| 0.015 | 0.0158844 |
| 0.0175 | 0.0200087 |
| 0.02 | 0.024725 |
| 0.0225 | 0.0301182 |
| 0.025 | 0.0362855 |
| 0.0275 | 0.043338 |
| 0.03 | 0.0514027 |
| 0.0325 | 0.0606249 |
| 0.035 | 0.0711708 |
| 0.0375 | 0.0832303 |
| 0.04 | 0.0970208 |
| 0.0425 | 0.112791 |
| 0.045 | 0.130824 |
| 0.0475 | 0.151445 |
| 0.05 | 0.175026 |
| 0.0525 | 0.201992 |
| 0.055 | 0.232828 |
| 0.0575 | 0.26809 |
| 0.06 | 0.308414 |
| 0.0625 | 0.354524 |
| 0.065 | 0.407253 |
| 0.0675 | 0.46755 |
| 0.07 | 0.536502 |
| 0.0725 | 0.61535 |
| 0.075 | 0.705514 |
| 0.0775 | 0.808621 |
| 0.08 | 0.926526 |

| Strain (mm/mm) | CCL3_Asc |
| --- | --- |
| 0 | 0 |
| 0.0025 | 0.00238689 |
| 0.005 | 0.00526029 |
| 0.0075 | 0.00871938 |
| 0.01 | 0.0128835 |
| 0.0125 | 0.0178965 |
| 0.015 | 0.0239312 |
| 0.0175 | 0.031196 |
| 0.02 | 0.0399416 |
| 0.0225 | 0.0504698 |
| 0.025 | 0.0631439 |
| 0.0275 | 0.0784014 |
| 0.03 | 0.0967689 |
| 0.0325 | 0.11888 |
| 0.035 | 0.145499 |
| 0.0375 | 0.177542 |
| 0.04 | 0.216118 |
| 0.0425 | 0.262556 |
| 0.045 | 0.31846 |

Stress (MPa)

| Strain (mm/mm) | CCL4_Desc |
| --- | --- |
| 0 | 0 |
| 0.0025 | 0.00191636 |
| 0.005 | 0.00412038 |
| 0.0075 | 0.00665525 |
| 0.01 | 0.00957062 |
| 0.0125 | 0.0129236 |
| 0.015 | 0.01678 |
| 0.0175 | 0.0212152 |
| 0.02 | 0.0263161 |
| 0.0225 | 0.0321828 |
| 0.025 | 0.0389302 |
| 0.0275 | 0.0466903 |
| 0.03 | 0.0556154 |
| 0.0325 | 0.0658802 |
| 0.035 | 0.0776859 |
| 0.0375 | 0.0912637 |
| 0.04 | 0.10688 |
| 0.0425 | 0.12484 |
| 0.045 | 0.145496 |
| 0.0475 | 0.169253 |
| 0.05 | 0.196576 |
| 0.0525 | 0.228 |
| 0.055 | 0.264142 |
| 0.0575 | 0.305709 |
| 0.06 | 0.353515 |
| 0.0625 | 0.408498 |
| 0.065 | 0.471734 |
| 0.0675 | 0.544463 |
| 0.07 | 0.628108 |

| Strain (mm/mm) | CCL4_Asc |
| --- | --- |
| 0 | 0 |
| 0.0025 | 0.00146838 |
| 0.005 | 0.00314704 |
| 0.0075 | 0.00506607 |
| 0.01 | 0.00725991 |
| 0.0125 | 0.0097679 |
| 0.015 | 0.012635 |
| 0.0175 | 0.0159127 |
| 0.02 | 0.0196598 |
| 0.0225 | 0.0239434 |
| 0.025 | 0.0288405 |
| 0.0275 | 0.0344387 |
| 0.03 | 0.0408387 |
| 0.0325 | 0.0481551 |
| 0.035 | 0.0565192 |
| 0.0375 | 0.0660811 |
| 0.04 | 0.0770121 |
| 0.0425 | 0.0895085 |
| 0.045 | 0.103794 |
| 0.0475 | 0.120126 |
| 0.05 | 0.138796 |
| 0.0525 | 0.16014 |
| 0.055 | 0.18454 |
| 0.0575 | 0.212434 |
| 0.06 | 0.244323 |
| 0.0625 | 0.280778 |
| 0.065 | 0.322453 |
| 0.0675 | 0.370096 |
| 0.07 | 0.424562 |
| 0.0725 | 0.486827 |
| 0.075 | 0.558008 |
| 0.0775 | 0.639382 |
| 0.08 | 0.732408 |

Stress (MPa)

| Strain (mm/mm) | CCL5_Desc |
| --- | --- |
| 0 | 0 |
| 0.0025 | 0.00143387 |
| 0.005 | 0.00304292 |
| 0.0075 | 0.00484856 |
| 0.01 | 0.00687481 |
| 0.0125 | 0.00914862 |
| 0.015 | 0.0117002 |
| 0.0175 | 0.0145636 |
| 0.02 | 0.0177768 |
| 0.0225 | 0.0213826 |
| 0.025 | 0.025429 |
| 0.0275 | 0.0299697 |
| 0.03 | 0.0350652 |
| 0.0325 | 0.0407832 |
| 0.035 | 0.0471999 |
| 0.0375 | 0.0544005 |
| 0.04 | 0.0624809 |
| 0.0425 | 0.0715485 |
| 0.045 | 0.081724 |
| 0.0475 | 0.0931427 |
| 0.05 | 0.105956 |
| 0.0525 | 0.120336 |
| 0.055 | 0.136472 |
| 0.0575 | 0.15458 |
| 0.06 | 0.1749 |
| 0.0625 | 0.197702 |
| 0.065 | 0.223291 |
| 0.0675 | 0.252006 |
| 0.07 | 0.284229 |
| 0.0725 | 0.32039 |
| 0.075 | 0.360968 |
| 0.0775 | 0.406504 |
| 0.08 | 0.457604 |
| 0.0825 | 0.514946 |
| 0.085 | 0.579295 |
| 0.0875 | 0.651506 |
| 0.09 | 0.732539 |

| Strain (mm/mm) | CCL5_Asc |
| --- | --- |
| 0 | 0 |
| 0.0025 | 0.00140042 |
| 0.005 | 0.00299541 |
| 0.0075 | 0.00481199 |
| 0.01 | 0.00688096 |
| 0.0125 | 0.00923737 |
| 0.015 | 0.0119212 |
| 0.0175 | 0.0149778 |
| 0.02 | 0.0184591 |
| 0.0225 | 0.0224241 |
| 0.025 | 0.02694 |
| 0.0275 | 0.0320832 |
| 0.03 | 0.037941 |
| 0.0325 | 0.0446127 |
| 0.035 | 0.0522112 |
| 0.0375 | 0.0608655 |
| 0.04 | 0.0707221 |
| 0.0425 | 0.081948 |
| 0.045 | 0.0947337 |
| 0.0475 | 0.109296 |
| 0.05 | 0.125881 |
| 0.0525 | 0.14477 |
| 0.055 | 0.166284 |
| 0.0575 | 0.190786 |
| 0.06 | 0.218693 |
| 0.0625 | 0.250477 |
| 0.065 | 0.286676 |
| 0.0675 | 0.327905 |
| 0.07 | 0.374862 |
| 0.0725 | 0.428343 |
| 0.075 | 0.489254 |
| 0.0775 | 0.558627 |
| 0.08 | 0.637639 |

Stress (MPa)

| Strain (mm/mm) | S6_Desc |
| --- | --- |
| 0 | 0 |
| 0.0025 | 0.00148807 |
| 0.005 | 0.00318705 |
| 0.0075 | 0.00512683 |
| 0.01 | 0.00734154 |
| 0.0125 | 0.00987016 |
| 0.015 | 0.0127572 |
| 0.0175 | 0.0160534 |
| 0.02 | 0.0198167 |
| 0.0225 | 0.0241135 |
| 0.025 | 0.0290193 |
| 0.0275 | 0.0346204 |
| 0.03 | 0.0410153 |
| 0.0325 | 0.0483166 |
| 0.035 | 0.0566528 |
| 0.0375 | 0.0661705 |
| 0.04 | 0.0770372 |
| 0.0425 | 0.089444 |
| 0.045 | 0.103609 |
| 0.0475 | 0.119782 |
| 0.05 | 0.138248 |
| 0.0525 | 0.15933 |
| 0.055 | 0.183401 |
| 0.0575 | 0.210883 |
| 0.06 | 0.24226 |
| 0.0625 | 0.278085 |
| 0.065 | 0.318987 |
| 0.0675 | 0.365686 |
| 0.07 | 0.419004 |
| 0.0725 | 0.479879 |
| 0.075 | 0.549382 |
| 0.0775 | 0.628737 |

| Strain (mm/mm) | CCL6_Asc |
| --- | --- |
| 0 | 0 |
| 0.0025 | 0.00172174 |
| 0.005 | 0.00371078 |
| 0.0075 | 0.00600862 |
| 0.01 | 0.00866319 |
| 0.0125 | 0.0117299 |
| 0.015 | 0.0152727 |
| 0.0175 | 0.0193655 |
| 0.02 | 0.0240937 |
| 0.0225 | 0.0295559 |
| 0.025 | 0.0358662 |
| 0.0275 | 0.0431561 |
| 0.03 | 0.0515778 |
| 0.0325 | 0.0613069 |
| 0.035 | 0.0725465 |
| 0.0375 | 0.085531 |
| 0.04 | 0.100531 |
| 0.0425 | 0.11786 |
| 0.045 | 0.13788 |
| 0.0475 | 0.161007 |
| 0.05 | 0.187725 |
| 0.0525 | 0.218591 |
| 0.055 | 0.254249 |
| 0.0575 | 0.295442 |
| 0.06 | 0.343031 |
| 0.0625 | 0.398008 |

**Fig. S4.** The stress-strain characteristics of individual canine cranial cruciate ligaments (CCL) during the ascending (Asc) and descending (Desc) tests at 1%/min strain rate.

Stress (MPa)

| Strain (mm/mm) | CCL1_Desc |
| --- | --- |
| 0 | 0 |
| 0.0025 | 0.000964313 |
| 0.005 | 0.00199706 |
| 0.0075 | 0.0031031 |
| 0.01 | 0.00428763 |
| 0.0125 | 0.00555623 |
| 0.015 | 0.00691485 |
| 0.0175 | 0.00836989 |
| 0.02 | 0.00992819 |
| 0.0225 | 0.0115971 |
| 0.025 | 0.0133844 |
| 0.0275 | 0.0152986 |
| 0.03 | 0.0173486 |
| 0.0325 | 0.0195441 |
| 0.035 | 0.0218954 |
| 0.0375 | 0.0244135 |
| 0.04 | 0.0271104 |
| 0.0425 | 0.0299987 |
| 0.045 | 0.0330919 |
| 0.0475 | 0.0364046 |
| 0.05 | 0.0399525 |
| 0.0525 | 0.0437521 |
| 0.055 | 0.0478214 |
| 0.0575 | 0.0521794 |
| 0.06 | 0.0568468 |
| 0.0625 | 0.0618453 |
| 0.065 | 0.0671986 |
| 0.0675 | 0.0729319 |
| 0.07 | 0.0790719 |
| 0.0725 | 0.0856477 |
| 0.075 | 0.0926902 |
| 0.0775 | 0.100233 |
| 0.08 | 0.10831 |
| 0.0825 | 0.116961 |
| 0.085 | 0.126225 |
| 0.0875 | 0.136148 |
| 0.09 | 0.146774 |
| 0.0925 | 0.158154 |
| 0.095 | 0.170342 |
| 0.0975 | 0.183395 |
| 0.1 | 0.197375 |
| 0.1025 | 0.212346 |
| 0.105 | 0.22838 |
| 0.1075 | 0.245552 |
| 0.11 | 0.263942 |
| 0.1125 | 0.283638 |
| 0.115 | 0.304731 |
| 0.1175 | 0.327321 |
| 0.12 | 0.351515 |
| 0.1225 | 0.377425 |
| 0.125 | 0.405174 |
| 0.1275 | 0.434893 |
| 0.13 | 0.46672 |
| 0.1325 | 0.500806 |
| 0.135 | 0.537311 |
| 0.1375 | 0.576407 |
| 0.14 | 0.618277 |
| 0.1425 | 0.663119 |
| 0.145 | 0.711143 |
| 0.1475 | 0.762575 |
| 0.15 | 0.817657 |

| Strain (mm/mm) | CCL1_Asc |
| --- | --- |
| 0 | 0 |
| 0.0025 | 0.00375745 |
| 0.005 | 0.00803218 |
| 0.0075 | 0.0128954 |
| 0.01 | 0.0184282 |
| 0.0125 | 0.0247226 |
| 0.015 | 0.0318836 |
| 0.0175 | 0.0400304 |
| 0.02 | 0.0492988 |
| 0.0225 | 0.0598431 |
| 0.025 | 0.0718391 |
| 0.0275 | 0.0854866 |
| 0.03 | 0.101013 |
| 0.0325 | 0.118677 |
| 0.035 | 0.138772 |
| 0.0375 | 0.161634 |
| 0.04 | 0.187644 |
| 0.0425 | 0.217234 |
| 0.045 | 0.250898 |
| 0.0475 | 0.289196 |
| 0.05 | 0.332766 |
| 0.0525 | 0.382335 |
| 0.055 | 0.438729 |
| 0.0575 | 0.502885 |
| 0.06 | 0.575874 |
| 0.0625 | 0.658912 |

Stress (MPa)

| Strain (mm/mm) | CCL2_Desc |
| --- | --- |
| 0 | 0 |
| 0.0025 | 0.00053044 |
| 0.005 | 0.00122662 |
| 0.0075 | 0.00214033 |
| 0.01 | 0.00333953 |
| 0.0125 | 0.00491342 |
| 0.015 | 0.00697909 |
| 0.0175 | 0.0096902 |
| 0.02 | 0.0132484 |
| 0.0225 | 0.0179184 |
| 0.025 | 0.0240476 |
| 0.0275 | 0.0320918 |
| 0.03 | 0.0426496 |
| 0.0325 | 0.0565061 |
| 0.035 | 0.0746923 |
| 0.0375 | 0.0985608 |
| 0.04 | 0.129887 |
| 0.0425 | 0.171002 |
| 0.045 | 0.224963 |
| 0.0475 | 0.295784 |
| 0.05 | 0.388735 |
| 0.0525 | 0.510728 |
| 0.055 | 0.670839 |

| Strain (mm/mm) | CCL2_Asc |
| --- | --- |
| 0 | 0 |
| 0.0025 | 0.00246139 |
| 0.005 | 0.00543084 |
| 0.0075 | 0.0090132 |
| 0.01 | 0.013335 |
| 0.0125 | 0.0185489 |
| 0.015 | 0.0248389 |
| 0.0175 | 0.0324273 |
| 0.02 | 0.041582 |
| 0.0225 | 0.0526263 |
| 0.025 | 0.0659502 |
| 0.0275 | 0.0820243 |
| 0.03 | 0.101416 |
| 0.0325 | 0.124811 |
| 0.035 | 0.153035 |
| 0.0375 | 0.187084 |
| 0.04 | 0.228161 |
| 0.0425 | 0.277717 |
| 0.045 | 0.337501 |
| 0.0475 | 0.409626 |
| 0.05 | 0.496638 |
| 0.0525 | 0.601611 |

Stress (MPa)

| Strain (mm/mm) | CCL3_Desc |
| --- | --- |
| 0 | 0 |
| 0.0025 | 0.0021677 |
| 0.005 | 0.00465231 |
| 0.0075 | 0.00750018 |
| 0.01 | 0.0107644 |
| 0.0125 | 0.0145059 |
| 0.015 | 0.0187943 |
| 0.0175 | 0.0237097 |
| 0.02 | 0.0293438 |
| 0.0225 | 0.0358016 |
| 0.025 | 0.0432035 |
| 0.0275 | 0.0516875 |
| 0.03 | 0.0614119 |
| 0.0325 | 0.0725581 |
| 0.035 | 0.0853337 |
| 0.0375 | 0.0999773 |
| 0.04 | 0.116762 |
| 0.0425 | 0.136 |
| 0.045 | 0.158051 |
| 0.0475 | 0.183326 |
| 0.05 | 0.212295 |
| 0.0525 | 0.245501 |
| 0.055 | 0.283561 |
| 0.0575 | 0.327185 |
| 0.06 | 0.377187 |
| 0.0625 | 0.4345 |
| 0.065 | 0.500191 |
| 0.0675 | 0.575487 |
| 0.07 | 0.661791 |
| 0.0725 | 0.760713 |
| 0.075 | 0.874097 |

| Strain (mm/mm) | CCL3_Asc |
| --- | --- |
| 0 | 0 |
| 0.0025 | 0.00241894 |
| 0.005 | 0.00534862 |
| 0.0075 | 0.00889688 |
| 0.01 | 0.0131943 |
| 0.0125 | 0.0183992 |
| 0.015 | 0.024703 |
| 0.0175 | 0.0323378 |
| 0.02 | 0.0415847 |
| 0.0225 | 0.052784 |
| 0.025 | 0.066348 |
| 0.0275 | 0.0827759 |
| 0.03 | 0.102672 |
| 0.0325 | 0.12677 |
| 0.035 | 0.155956 |
| 0.0375 | 0.191304 |
| 0.04 | 0.234116 |
| 0.0425 | 0.285967 |

Stress (MPa)

| Strain (mm/mm) | CCL4_Desc |
| --- | --- |
| 0 | 0 |
| 0.0025 | 0.00210369 |
| 0.005 | 0.00452536 |
| 0.0075 | 0.00731306 |
| 0.01 | 0.0105221 |
| 0.0125 | 0.0142162 |
| 0.015 | 0.0184687 |
| 0.0175 | 0.0233639 |
| 0.02 | 0.028999 |
| 0.0225 | 0.0354858 |
| 0.025 | 0.0429532 |
| 0.0275 | 0.0515492 |
| 0.03 | 0.0614444 |
| 0.0325 | 0.0728354 |
| 0.035 | 0.085948 |
| 0.0375 | 0.101043 |
| 0.04 | 0.118419 |
| 0.0425 | 0.138421 |
| 0.045 | 0.161447 |
| 0.0475 | 0.187953 |
| 0.05 | 0.218466 |
| 0.0525 | 0.253591 |
| 0.055 | 0.294024 |
| 0.0575 | 0.340569 |
| 0.06 | 0.39415 |
| 0.0625 | 0.455828 |
| 0.065 | 0.52683 |
| 0.0675 | 0.608563 |

| Strain (mm/mm) | CCL4_Asc |
| --- | --- |
| 0 | 0 |
| 0.0025 | 0.00161547 |
| 0.005 | 0.00346773 |
| 0.0075 | 0.00559152 |
| 0.01 | 0.00802662 |
| 0.0125 | 0.0108187 |
| 0.015 | 0.01402 |
| 0.0175 | 0.0176906 |
| 0.02 | 0.0218992 |
| 0.0225 | 0.0267248 |
| 0.025 | 0.0322577 |
| 0.0275 | 0.0386017 |
| 0.03 | 0.0458756 |
| 0.0325 | 0.0542158 |
| 0.035 | 0.0637785 |
| 0.0375 | 0.0747429 |
| 0.04 | 0.0873146 |
| 0.0425 | 0.101729 |
| 0.045 | 0.118257 |
| 0.0475 | 0.137207 |
| 0.05 | 0.158935 |
| 0.0525 | 0.183848 |
| 0.055 | 0.212412 |
| 0.0575 | 0.245164 |
| 0.06 | 0.282717 |
| 0.0625 | 0.325775 |
| 0.065 | 0.375145 |
| 0.0675 | 0.431751 |
| 0.07 | 0.496654 |
| 0.0725 | 0.571072 |
| 0.075 | 0.656398 |

Stress (MPa)

| Strain (mm/mm) | CCL5_Desc |
| --- | --- |
| 0 | 0 |
| 0.0025 | 0.00166808 |
| 0.005 | 0.00354394 |
| 0.0075 | 0.00565346 |
| 0.01 | 0.00802573 |
| 0.0125 | 0.0106935 |
| 0.015 | 0.0136936 |
| 0.0175 | 0.0170673 |
| 0.02 | 0.0208613 |
| 0.0225 | 0.0251279 |
| 0.025 | 0.0299259 |
| 0.0275 | 0.0353216 |
| 0.03 | 0.0413893 |
| 0.0325 | 0.0482129 |
| 0.035 | 0.0558863 |
| 0.0375 | 0.0645156 |
| 0.04 | 0.0742198 |
| 0.0425 | 0.0851327 |
| 0.045 | 0.0974049 |
| 0.0475 | 0.111206 |
| 0.05 | 0.126726 |
| 0.0525 | 0.144179 |
| 0.055 | 0.163806 |
| 0.0575 | 0.185878 |
| 0.06 | 0.210699 |
| 0.0625 | 0.238611 |
| 0.065 | 0.270001 |
| 0.0675 | 0.3053 |
| 0.07 | 0.344997 |
| 0.0725 | 0.389638 |
| 0.075 | 0.439839 |
| 0.0775 | 0.496294 |
| 0.08 | 0.55978 |
| 0.0825 | 0.631175 |
| 0.085 | 0.711462 |
| 0.0875 | 0.80175 |

| Strain (mm/mm) | CCL5_Asc |
| --- | --- |
| 0 | 0 |
| 0.0025 | 0.00150253 |
| 0.005 | 0.00322118 |
| 0.0075 | 0.00518703 |
| 0.01 | 0.00743565 |
| 0.0125 | 0.0100077 |
| 0.015 | 0.0129497 |
| 0.0175 | 0.0163149 |
| 0.02 | 0.0201641 |
| 0.0225 | 0.024567 |
| 0.025 | 0.0296032 |
| 0.0275 | 0.0353638 |
| 0.03 | 0.041953 |
| 0.0325 | 0.0494899 |
| 0.035 | 0.0581109 |
| 0.0375 | 0.067972 |
| 0.04 | 0.0792515 |
| 0.0425 | 0.0921534 |
| 0.045 | 0.106911 |
| 0.0475 | 0.123791 |
| 0.05 | 0.1431 |
| 0.0525 | 0.165185 |
| 0.055 | 0.190448 |
| 0.0575 | 0.219344 |
| 0.06 | 0.252396 |
| 0.0625 | 0.290203 |
| 0.065 | 0.333448 |
| 0.0675 | 0.382913 |
| 0.07 | 0.439492 |
| 0.0725 | 0.50421 |
| 0.075 | 0.578237 |

Stress (MPa)

| Strain (mm/mm) | S6_Desc |
| --- | --- |
| 0 | 0 |
| 0.0025 | 0.00146638 |
| 0.005 | 0.0031418 |
| 0.0075 | 0.00505607 |
| 0.01 | 0.00724324 |
| 0.0125 | 0.00974221 |
| 0.015 | 0.0125974 |
| 0.0175 | 0.0158597 |
| 0.02 | 0.0195871 |
| 0.0225 | 0.0238458 |
| 0.025 | 0.0287116 |
| 0.0275 | 0.0342711 |
| 0.03 | 0.0406232 |
| 0.0325 | 0.0478809 |
| 0.035 | 0.0561732 |
| 0.0375 | 0.0656476 |
| 0.04 | 0.0764727 |
| 0.0425 | 0.0888411 |
| 0.045 | 0.102973 |
| 0.0475 | 0.119119 |
| 0.05 | 0.137567 |
| 0.0525 | 0.158645 |
| 0.055 | 0.182728 |
| 0.0575 | 0.210244 |
| 0.06 | 0.241683 |
| 0.0625 | 0.277604 |
| 0.065 | 0.318645 |
| 0.0675 | 0.365538 |
| 0.07 | 0.419116 |
| 0.0725 | 0.480331 |
| 0.075 | 0.550274 |
| 0.0775 | 0.630188 |

| Strain (mm/mm) | CCL6_Asc |
| --- | --- |
| 0 | 0 |
| 0.0025 | 0.00170073 |
| 0.005 | 0.00367418 |
| 0.0075 | 0.00596408 |
| 0.01 | 0.00862118 |
| 0.0125 | 0.0117043 |
| 0.015 | 0.0152819 |
| 0.0175 | 0.0194332 |
| 0.02 | 0.0242501 |
| 0.0225 | 0.0298394 |
| 0.025 | 0.036325 |
| 0.0275 | 0.0438506 |
| 0.03 | 0.052583 |
| 0.0325 | 0.0627157 |
| 0.035 | 0.0744731 |
| 0.0375 | 0.0881159 |
| 0.04 | 0.103946 |
| 0.0425 | 0.122315 |
| 0.045 | 0.14363 |
| 0.0475 | 0.168362 |
| 0.05 | 0.197061 |
| 0.0525 | 0.230361 |
| 0.055 | 0.269001 |
| 0.0575 | 0.313837 |
| 0.06 | 0.365863 |
| 0.0625 | 0.426231 |

**Fig. S5.** The stress-strain characteristics of individual canine cranial cruciate ligaments (CCL) during the ascending (Asc) and descending (Desc) tests at 10%/min strain rate.
